# An Extensible Vector Toolkit and Parts Library for Advanced Engineering of Plant Genomes

**DOI:** 10.1101/2022.10.15.511792

**Authors:** James C. Chamness, Jitesh Kumar, Anna J. Cruz, Elissa Rhuby, Mason J. Holum, Jon P. Cody, Redeat Tibebu, Maria Elena Gamo, Colby G. Starker, Feng Zhang, Daniel F. Voytas

## Abstract

Plant biotechnology is rife with new advances in transformation and genome engineering techniques. A common requirement for delivery and coordinated expression in plant cells, however, places the design and assembly of transformation constructs at a crucial juncture as desired reagent suites grow more complex. Modular cloning principles have simplified some aspects of vector design, yet many important components remain unavailable or poorly adapted for rapid implementation in biotechnology research. Here, we describe a universal Golden Gate cloning toolkit for vector construction. The toolkit chassis is compatible with the widely accepted Phytobrick standard for genetic parts, and supports assembly of arbitrarily complex T-DNAs through improved capacity, positional flexibility, and extensibility in comparison to extant kits. We also provision a substantial library of newly adapted Phytobricks, including regulatory elements for monocot and dicot gene expression, and coding sequences for genes of interest such as reporters, developmental regulators, and site-specific recombinases. Finally, we use a series of dual luciferase assays to measure contributions to expression from promoters, terminators, and from cross-cassette interactions attributable to enhancer elements in certain promoters. Taken together, these publicly available cloning resources can greatly accelerate the testing and deployment of new tools for plant engineering.

## Introduction

Transformation is a cornerstone of plant biotechnology, underpinning the majority of practical genome engineering techniques and approaches to plant synthetic biology. Transformation constructs thus play a key role in advancing biotechnology efforts in plants. The inefficiencies of DNA delivery and regeneration of transformed cells dictate many features of vector design prior to consideration of downstream applications. The use of Agrobacterium for transformation, a method preferred for simpler integration patterns in comparison to physical delivery methods^1,2^, imposes a unique requirement to optimize vector behavior across three separate hosts: *E. coli* for cloning, Agrobacterium for infection, and the plant subject. Most T-DNA vectors include markers for chemical or visual selection of transformed cells and, increasingly, it is desirable to include developmental regulator genes (DRs) to stimulate growth in the primary explants, which can dramatically improve regeneration of transgenic or edited plants. However, DRs must often be paired with tissue-specific promoters or other expression control systems to avoid deleterious effects of ectopic expression^3^.

The widening array of tools for DR-mediated transformation, genome engineering, and synthetic biology all contribute to a trending increase in vector complexity. This reflects an increasing number of genes on each construct, but also requirements for their specific and orthogonal expression. Considering constructs as genetic programs played out through plant cell lineages, the most important factor under experimenter control for achieving the desired transgene effects is the choice of *cis*-regulatory elements governing their expression. Varying the promoter driving a selectable marker impacts transformation efficiency^4,5^; the promoters driving editing reagents similarly affect editing frequencies, particularly for targeting germline cells that can give rise to heritable edits^6–8^, and synthetic promoters underpin the tightly regulated expression crucial for genetic circuits and precise metabolic engineering^9^. Variation in terminators also impacts gene expression, though the extent remains unclear^10,11^. Beyond promoter and terminator selection, other, less intuitive factors such as cassette orientation, homology-dependent silencing, and cross-cassette interactions may all influence transgene expression, and merit close consideration at the point of construct design^12,13^.

The increasing scope of genetic elements used in transformation constructs have motivated efforts to standardize cloning methods, and facilitate reuse and exchange of parts between groups. The most successful standard is a modular cloning scheme based on Golden Gate assembly with type IIS restriction enzymes, first described in the “MoClo” vector toolkit from the Marillonet group in 2011^14^. This standard subdivides a transcriptional unit into a maximum of ten functional parts, each defined by unique, flanking 4-base overhangs. Discrete parts are carried on “Level 0” plasmids, alternately referred to in the literature as MoClo parts or, more recently, as Phytobricks (to disambiguate from a higher-level vector assembly scheme described in the same report). Any set of Level 0 parts with contiguous overhangs can be assembled into a “Level 1” destination vector, via BsaI Golden Gate assembly, to build a Level 1 module constituting a complete transcriptional unit (TU). The Phytobrick standard has received widespread adoption, first among plant synthetic biologists but increasingly in the broader plant research community^15^. Multiple Phytobrick collections have been published and made available through public vector repositories such as Addgene, including various coding sequences, *cis*-regulatory elements, gene editing tools, and other elements of interest for plant biotechnology^16–20^.

While the Phytobrick interface for genetic parts is now broadly accepted, there is greater divergence among strategies for higher-level assembly of Level 1 modules into multigenic transformation constructs. Several cloning toolkits have been described which are Phytobrick-compatible but otherwise non-interoperable. The original MoClo kit^14^ follows a hierarchical model, wherein Level 1 modules with single TUs are assembled into “Level 2” vectors, typically binary vectors, to create transformation constructs. The GoldenBraid^21^, Mobius^22^, and Loop^23^ kits all follow a looping model, wherein at least two classes of modules may be alternately assembled together by cycling between different bacterial selections and restriction enzymes, concatenating multiple TUs into single cloning units at each step. By using specialized destination vectors with binary vector components for one or either module class, any cloning unit can be entered into a T-DNA to create a transformation construct. These looping schemes permit theoretically infinite expansion, limited only by the efficiency of large plasmid assembly and stability in *E. coli*. An extension to the MoClo kit introduced similar looping capability, using analogous intermediate layers for module concatenation.^24^ For an illustrated comparison of assembly schemes, and a complete description of the Phytobrick standard, see Cai, Lopez & Patron (2020)^25^.

In this work, we describe a set of publicly available resources for modular cloning: a diverse library of genetic parts for plant engineering, newly adapted to the Phytobrick standard and thus compatible with all leading vector toolkits; a set of binary vector backbones, incorporating improvements from previous works; and a novel cloning framework for higher-level assembly, which offers a greater compromise of capacity, flexibility, and extensibility in comparison to previous toolkits. We also present experiments investigating the impact of *cis*-regulatory elements on transgene expression, illustrating how our cloning framework enables rapid testing of components for plant transformation.

## Materials & Methods

### Construction of modular cloning toolkit

A bidirectional cloning block with a chloramphenicol (cam) resistance marker and *ccdB* was first constructed by modifying a version present in vectors from Cermak *et al.* (2017)^26^ to domesticate for BsaI and Esp3I sites and remove unused flanking sequences for Gateway assembly. Various iterations of this block were used to construct the toolkit vectors by adding different Golden Gate enzymes and overhangs with PCR primers, and assembling these together with amplified backbone fragments. The Level 0 destination vector backbones were amplified from the analogous *lacZα* versions in Engler *et al.* (2014)^16^. The Level 1 destination vector backbones were amplified from the carbenicillin (carb) modules described in Cermak *et al.* (2017)^26^. The Golden Gate extension modules were assembled by replacing annealed oligo linkers into the corresponding Level 1 destination vectors, using PaqCI for traditional digestion and ligation. The destination donor modules were assembled by blunt ligation of amplicons of the primary cloning block into a domesticated version of the PJET cloning vector digested with EcoRV. The Level 2 destination vectors were constructed using both Golden Gate and Gibson cloning of fragments from multiple sources. The common pUC19 origin and kanamycin (kan) marker used in all the pMIN vectors was sourced from pLSU4mg10 (Addgene #126079)^27^. The pVS1 origin was sourced from the pTRANS vectors^26^, and used to assemble pMIN-VS1 following the design pattern of the pLSU vectors^28^. The pRi origin was synthesized from TWIST Bioscience, using a demarcation for the ends of a minimal replicon described in Ye *et al.* (2011)^29^. All vectors carrying *ccdB* were propagated in the resistant *E. coli* strain DB3.1 (ThermoFisher, discontinued; available from Abbexa, abx098865), and all other vectors were propagated in either NEB10*β* Competent or NEB Stable Competent cells (NEB C3019I, C3040I).

### Construction of Phytobrick parts library

Level 0 parts were synthesized as linear fragments from IDT or TWIST Bioscience, or amplified from various sources, including pre-existing plasmids and genomic DNA from maize, sorghum, rice, tomato, potato, and tobacco; a complete list of sources is provided in Table S1. Modifications for domestication were introduced as design changes prior to synthesis, or with PCR mutagenesis during initial assembly. All fragments were entered into Level 0 destination vectors via BbsI Golden Gate assembly.

### Dual-luciferase assays

Luciferase activity was measured with the Dual-Luciferase Reporter Assay kit (Promega E1960). For the leaf infiltration experiments, constructs were prepared in Agrobacterium strain GV3101 with the pMP90 helper plasmid and grown as overnight cultures. Cultures were resuspended in infiltration buffer (5 g/L glucose, 500mM MES), adjusted to an OD600 of 0.8, and infiltrated into leaves of approximately 1-month old *Nicotiana benthamiana* plants. At least 5 biological (infiltration) replicates were prepared for each vector. At 5 days after infiltration, 1cm discs were collected from Ruby-positive leaf tissue, flash frozen in liquid nitrogen, ground to powder with steel beads, and resuspended in 200*μ*L passive lysis buffer. Samples were scored using 20*μ*L aliquots loaded into a 96-well plate and run on a GloMax Explorer Multimode Plate Reader (Promega GM3500) using the kit-provided reagents and protocol, returning Relative Luminescence Unit (RLU) values for each sample for firefly and Renilla activity. The level of background signal was determined to be ≤ 150 RLUs for both luciferases, based on replicates with non-infiltated, wild-type leaf tissue, while signal in infiltrated leaves typically ranged from 10^3^ - 10^5^. For the promoter and terminator comparisons, a small number of samples were excluded from analysis due to measured luminescence values indistinguishable from background level; we inferred this was due to poor infection, as not all infiltrated leaves produced strong Ruby signal. For the remaining samples, we performed background subtraction, then calculated the ratio of firefly RLUs to Renilla RLUs to produce a metric for relative expression. Datapoints were then scaled by the mean FLuc/RLuc ratio among the samples for an internal control construct in each experiment: *Nos∷Firefly∷Nos* for the promoter group, and *2x35S∷TMV*Ω*∷Firefly∷Nos* for the terminator group, to measure the fold-change in expression versus these controls.

For the monocot promoter experiments in Setaria, leaves from approximately 3 week-old seedlings were used for protoplast isolation and transformation following the method of Weiss *et al.* (2020)^30^ with slight modifications. Briefly, three individual samples of 200K cells were incubated with 15 *μ*g of plasmid in 20% polyethylene glycol (PEG) solution, pH 5.7. Transfected cells were washed twice and incubated in W5 buffer (2mM MES, pH 5.7, 150mM NaCl, 125mM CaCl_2_, 5mM KCl) in the dark at room temperature for 36 hrs. Transfection efficiency was monitored through the expression of a GFP reporter gene. Samples were collected by centrifugation, resuspended in 100*μ*L passive lysis buffer, then scored for luciferase activity with the same plate reader assay. Based on 3 replicates transformed with a GFP control construct alone, the level of background signal was determined to be ≤ 200 RLUs for both luciferases, while signal in the dual-luciferase samples ranged from 10^2^ - 10^6^. Background subtraction was performed in the same fashion as the leaf infiltration samples, and all samples were retained in the final analysis. Due to the low signal from *Nos* and *2x35S* promoters, we chose to normalize fold-change expression to *CmYLCV∷Firefly∷Nos*, using the same scaling method as in the leaf infiltration experiments.

RNA-seq data for comparison with promoter expression results were drawn from the Tomato Functional Genomics Database, TFGP (http://ted.bti.cornell.edu/); The Arabidopsis Information Resource, TAIR (https://www.arabidopsis.org/index.jsp); MaizeGDB (https://www.maizegdb.org/); and the Rice Genome Annotation Project (http://rice.uga.edu/index.shtml).

For the promoter trans-activation experiments, we did not filter datapoints by signal level: all Renilla values were well above background, indicating effective delivery, and the majority of firefly values were, as expected, at or below background level. No normalization was performed as there was no internal control construct.

Within each experimental panel, significant differences between a construct and the given control were assessed using pairwise *t*-tests on an ordinary least squares linear regression, with Simes-Hochberg correction of P-values to account for multiple testing.

## Results & Discussion

### Joint Modular Cloning (JMC) Toolkit

#### Hierarchical assembly framework

Despite the preponderance of published toolkits, several factors motivated us to develop a new framework for higher-level assembly. Design philosophies for previous toolkits broadly reflect a tradeoff between greater capacity and complexity versus simplicity and size of the base vector set, but practical differences also arise through choice of Golden Gate enzymes, bacterial selections and cloning markers, and in the availability of different binary vector backbones for assembling T-DNA constructs. With respect to design philosophy, we find compelling advantages to the hierarchical model followed by the MoClo kit. First, this provides superior capacity for multigenic assembly in a single cloning step: MoClo offers seven unique Level 1 modules; Mobius and Loop each provide four; GoldenBraid provides two, though an extension to the original kit has increased this to five^31^. Another, more subjective advantage is that the hierarchical assembly model can be easier to learn and reason about than the looping model; we find this to be especially true when introducing new researchers to modular cloning. A further downside to the looping implementation is that all module concatenation must take place prior to entry into binary vectors, in order for the T-DNA border sequences to flank the final payload; i.e., binary vectors cannot be directly modified, and adding pieces to a previous assembly may require reworking of intermediate modules and an extra final cloning step. The only published mechanism for direct addition to pre-assembled binary vectors is the end-linker system from the MoClo kit, which uses specialized terminal modules with a bacterial color marker (CRed or *lacZα*) and auxiliary Golden Gate sites (BsaI or Esp3I) to append additional modules through a dual-enzyme Golden Gate reaction. However, this system limits expansion to a fixed point at the end of a construct, and the use of color marker genes within the end linkers makes the intermediate vectors less attractive as possible transformation constructs, since inclusion of these markers in plant transformation is likely undesirable.

In designing a new framework, we sought to incorporate strengths from each of these systems: the single-step capacity and intuitive design paradigm of the MoClo kit, repeatable extension as in the looping systems, and direct addition to binary vectors with arbitrary positional flexibility, which is not possible using any previous kit. The implementation simplifies selection of correct clones through exclusive use of lethal counterselections: all destination vectors, from Level 0 through Level 2, use *ccdB* as a negative marker, while all expansion vectors introduce a cam resistance marker (Figures S1, S2). This is in contrast to the color markers used in other major toolkits: MoClo, GoldenBraid, and Loop, for example, all use *lacZα*, which requires adding XGal and IPTG to bacterial plates for blue/white colony selection. Chloramphenicol provides an antibiotic selection, while the cytotoxic *ccdB* requires no additional reagents and prevents the formation of colonies which would otherwise appear as background.

The hierarchical model for the Joint Modular Cloning (JMC) toolkit is presented in Figure 1. New parts are generated with PCR or, increasingly, cost-effective synthesis (Figure 1a); once parts are entered into Level 0 destination vectors, all subsequent assemblies are PCR-free. Level 0 parts are assembled into Level 1 modules via BsaI Golden Gate, using the Phytobrick set of standard 4bp junctions with terminal overhangs J1 and J11 (Figure 1c). Not all internal overhangs are necessarily used in each assembly, as individual Phytobricks frequently span several of the more atomic part classes specified in the interface (Figure 1b). With carb selection, Level 1 modules are compatible with Phytobrick parts using either spectinomycin (spec) or cam selection. There are a total of 13 Level 1 module positions, for module chains of any length up to 7 (Figure S1). Modules are named by their chain position, with chain-terminating modules suffixed by “T”. The toolkit includes forward and reverse Phytobrick destination vectors for each module position to place cassettes in either orientation. For entry of components that cannot be domesticated for BsaI, the Level 1 destination vectors also include AscI and SacI sites for traditional restriction enzyme cloning (Figure S1).

**Figure 1:**
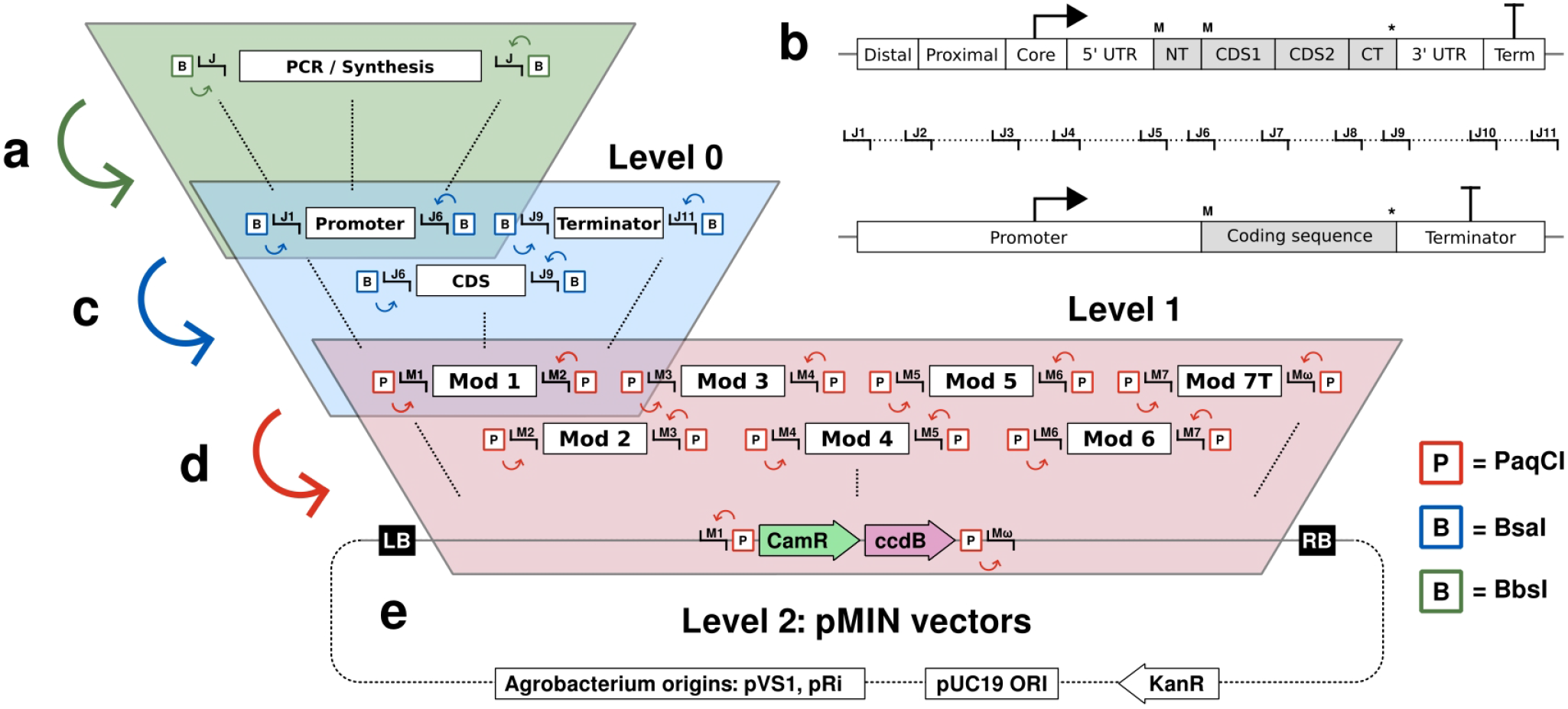
Hierarchical assembly of constructs from Phytobrick parts. **1a:** New parts are assembled into Level 0 destination vectors using BbsI with the Phytobrick (J1-J11) overhangs. **1b:** Phytobricks can occupy one of 10 atomic part classes, but commonly span simplified classes using J1, J6, J9, and J11. **1c:** Level 0 parts are assembled into Level 1 destination vectors using BsaI with J1 and J11. **1d:** Level 1 modules are assembled into Level 2 destination vectors using PaqCI with a new set of overhangs (M1-MΩ). **1e:** The pMIN vectors are Level 2 destination vectors, including a minimal backbone and 2 binary vector backbones.

Level 1 modules are assembled into Level 2 vectors with kan selection via PaqCI Golden Gate (Figure 1d). We selected PaqCI, an isoschizomer of AarI, for two reasons: a version of PaqCI has recently been optimized for Golden Gate assembly (NEB R0745S) and, as a rarer-cutting enzyme, it permits assembly of constructs from modules hosting components which cannot be domesticated for the more frequent cutters BsaI, Esp3I or BbsI. The overhangs used with PaqCI, termed M1-MΩ, were selected on the basis of simple heuristics for ligation fidelity: dissimilarity and absence of palindromes. However, the set includes multiple overhangs found to show high fidelity, and none found to show low fidelity, in recent work characterizing the behavior of 4bp overhangs in T4 ligations^32^. Experimentally, we have successfully used these overhangs to produce a high fraction of correct clones in assemblies up to and including all 7 modules.

It is worth noting that while the Phytobrick interface specifies part classes for the division of transcriptional units, the cloning mechanisms are agnostic to the abstract function of part sequences, excluding overhangs. Fragments with non-canonical function can thus be cloned using the same Level 0 or Level 1 vectors, to assemble modules or constructs with sequences such as insulators or recombinase recognition sites.

#### pMIN binary vectors for Agrobacterium transformation

For Level 2 assembly, we built 3 destination vectors: pMIN-0, pMIN-VS1, and pMIN-Ri. These vectors share kan selection, the pUC19 origin for *E. coli*, and have each been domesticated for all commonly used type IIS sites: PaqCI, BsaI, Esp3I, BbsI, and SapI. pMIN-0 is a minimal backbone with no binary vector components, suitable for physical transformation methods such as biolistics, PEG, and nanoparticle delivery. For Agrobacterium transformation, we constructed binary vectors from two different replicons: pVS1 and pRi. While some origins of replication, including the commonly used RK2 and pSa (P15A), function in both *E. coli* and Agrobacterium, we selected a dual-origin approach on the basis of plasmid stability and transformation performance. In Agrobacterium, plasmids based on both the pVS1 and pRi origins have shown superior stability in selection-free cultures to the RK2 origin^33,34^; the pSa replicon, used in the pGreen/pSoup system with replication proteins carried on a separate plasmid, was similarly found to be less stable when paired with either the RK2 or pVS1 origins, possibly due to a dosage imbalance of replication machinery attributable to differential plasmid copy-number^35,36^.

The pVS1 and pRi origins are considered high- and low-copy in Agrobacterium, respectively, with plasmids maintained at ~20 and ~2 copies per cell. Plasmid copy number has been found to influence transformation frequency and integration patterns, with pVS1 binary vectors leading to higher raw transformation frequencies, but pRi binary vectors leading to a greater fraction of single-copy events^37^. Either may be preferable, depending on experimental needs. Another consideration in selecting these origins was compatability with helper plasmids, since maintenance in a single strain requires compatible origins and selectable markers. Two recent reports describe ternary vector systems based on pVS1 helper plasmids: Anand *et al.* (2018) assessed the performance of different origin combinations for the T-DNA and helper plasmids, and found the best performing combination to be a pRi T-DNA vector and pVS1 helper plasmid: PHP71539, with gent selection^38^. Zhang *et al.* (2019) described similar, publicly available helper plasmids: pVS1-VIR2 and pRiA4-VIR, both with spec selection^36^. With kan selection, pMIN-Ri is compatible with both the PHP71539 and pVS1-VIR2 helper plasmids, and pMIN-VS1 with pRiA4-VIR; both are also compatible with the pMP90 helper plasmid with gent selection commonly used in GV3101.

Both pMIN-VS1 and pMIN-Ri are based on the smallest versions of these replicons described in the literature, for ease of manipulation *in vitro*. The pMIN-VS1 backbone is based on the pLSU vector architecture: this dispenses with the *bom* site, as triparental mating is now rarely used for plasmid introduction, and orients the two replicons of the binary plasmid for co-directional transcription, resulting in higher transformation efficiencies in *E. coli* and Agrobacterium, and higher plasmid yields in *E. coli* compared to traditional pVS1 vectors such as pCambia^28^. Our implementation further removes BsaI, Esp3I, and SapI sites from this backbone while installing a Golden Gate destination site within the T-DNA. The pMIN-Ri backbone is based on the minimal pRi replicon described in Ye *et al.* (2011), which dispenses with ~400bp downstream of RepC found to have no effect on propagation in Agrobacterium^29^. We further removed 2 PaqCI sites from the backbone, and introduced the same Golden Gate destination site as in pMIN-VS1. To the best of our knowledge, pMIN-Ri represents the first practical pRi-based modular cloning vector in the literature; the only other published Golden Gate backbone with the pRi origin is the MoClo toolkit vector pICH89921, a BIBAC vector based on the F-plasmid origin. This backbone is over 4kb larger than pMIN-Ri, and the single-copy F-plasmid origin requires specialized cultures to reach appreciable plasmid yields in *E. coli* preparations^39^. We have successfully conducted plant transformation with constructs built using both optimized backbones, using common Agrobacterium strains including AGL1, EHA105, GV3101, and LBA4404.

#### Modular expansion of binary vectors

To complement the hierarchical assembly framework, we developed a system for repeatable expansion of Level 2 constructs. Like the looping toolkits, this system enables theoretically infinite addition of modules; however, our system is based on direct addition to assembled transformation constructs, including binary vectors. The system consists of two complementary sets of vectors. The first are Level 1 expansion modules that function as any other Level 1 module in a Level 2 assembly with PaqCI, but instead of carrying transcriptional units, they contain ~30bp linkers with end-paired Esp3I or BsaI sites (Figure 2a). In contrast to the MoClo kit, these linkers lack cloning marker genes, so that the assembled Level 2 vectors are suitable as transformation constructs (Figure 2b). The second are a set of destination donor vectors that reintroduce a cloning marker and a new set of BsaI or PaqCI Golden Gate sites into the linker, creating a destination vector for subsequent assembly (Figure 2c). All the destination donors include cam resistance for positive selection of the new destination vector, and *ccdB* for negative selection in the downstream assembly, maintaining the toolkit invariant of lethal counterselection.

**Figure 2:**
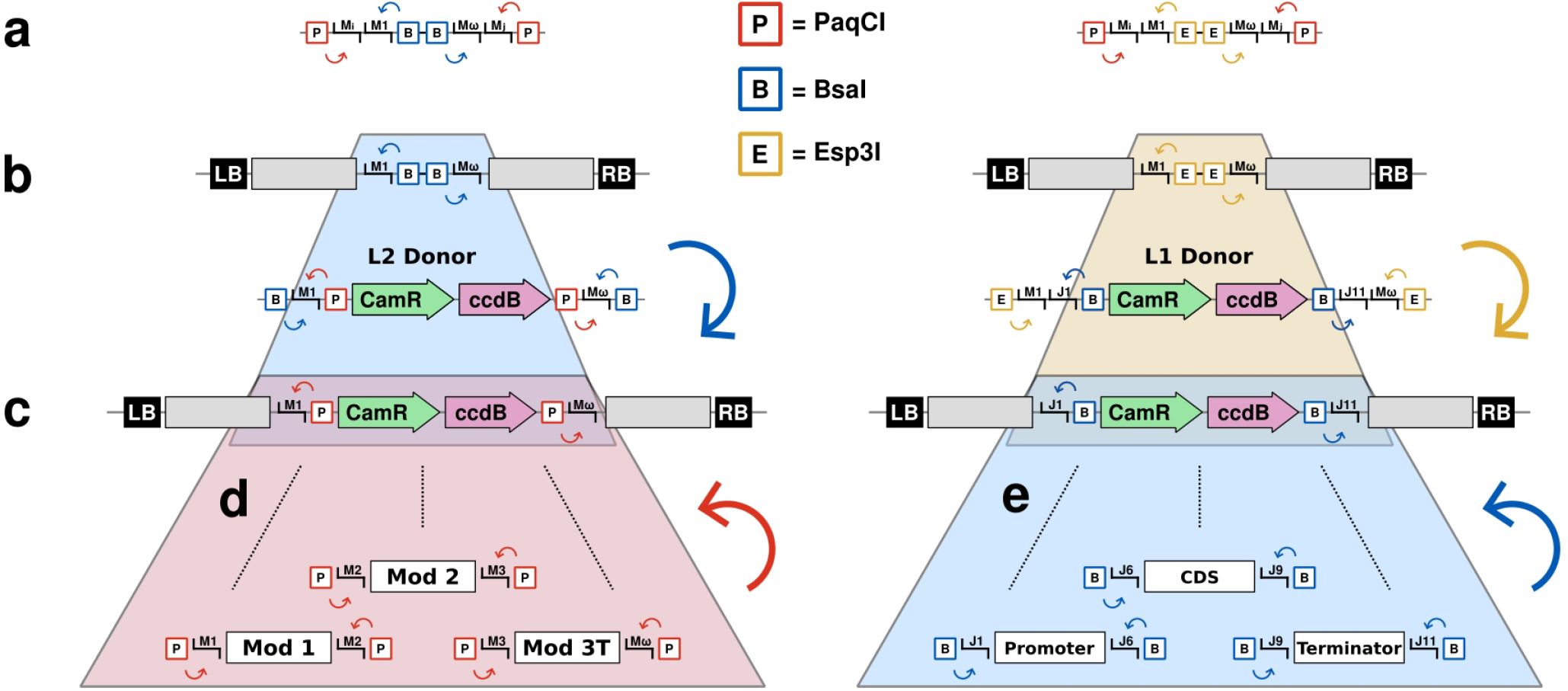
Modular expansion of binary vectors. **2a:** Expansion linkers are Level 1 modules at any position with end-paired Esp3I or BsaI sites. **2b:** Level 2 vectors with expansion linkers can be used directly as transformation constructs. **2c:** Destination donor vector restore a cloning marker and Golden Gate sites into an expansion linker. **2d:** Both BsaI and Esp3I linkers can be used to create Level 2 destination vectors. **2e:** Esp3I linkers can also be used to create Level 1 destination vectors.

This system can be used to create multiple types of destination vectors at any module position, and both Esp3I and BsaI linkers can be included in the same assembly to provide multiple orthogonal expansion points. Both linkers can be used with corresponding donors to create a “Level 2” destination point, with PaqCI and M1/MΩ (Figure 2d; Figure S2); repeated expansion is achieved by re-using linkers in the subsequent Level 2 assembly. The Esp3I linkers can also be used to create a “Level 1” destination point for Phytobrick parts with BsaI and J1/J11 (Figure 2e). Given forward and reverse orientation donor modules for both enzymes and destination classes, the system provides complete positional flexibility for assembly of arbitrarily complex constructs. The iterative assembly technique is particularly suited to parallelizing assembly of similar constructs with multiple common parts. For vector sets differing by a single cassette, the L1 destination donors can greatly simplify the number of required clones, as variant cassettes can be assembled from Phytobricks directly into a transformation backbone, requiring no Level 1 module intermediate; no comparable mechanism is provided in any other toolkit.

#### Phytobricks library

We have constructed a library of approximately a hundred genetic elements from the current literature. To adapt these elements to the Phytobrick standard and our toolkit, we domesticated all parts to remove BsaI and PaqCI recognition sites. Furthermore, we domesticated nearly all parts for Esp3I, BbsI, and SapI, to facilitate use with all other toolkits based on alternative Golden Gate enzymes. The majority of these parts have been functionally validated to ensure the SNPs introduced for domestication do not impact function. The largest set of parts are *cis*-regulatory elements, including viral, constitutive, tissue-specific, and inducible promoters for expression in both monocots and dicots; terminators from monocots and dicots; and genetic insulator sequences that attenuate enhancer interactions between neighboring cassettes^40–42^. We sought to provide a sufficient number of characterized elements to avoid repetition in multigenic constructs, as this increases the chances of transgene silencing in plant hosts^12^, and also the likelihood of recombination in bacterial hosts; we have observed recombination-mediated collapse of large T-DNAs with repeated elements, even in specialized cell lines such as NEB Stable *E. coli* and the *recA-* C58 derivative, AGL1^43^.

Library coding sequences include a variety of selectable marker and reporter genes. These include 2A fusions of markers and reporters to reduce cassette number requirements; non-traditional reporters such as AmCyan, an alternative to green and red fluorophores competing with autofluoresence in plant tissues; and reporters based on engineering metabolic pathways, including a fungal luciferase pathway and the Ruby reporter for betalain synthesis^44,45^. For DR transformation, we provide a set of the best-described genes, including *Baby boom* and *Wuschel2* from maize, a chimeric *GRF-GIF* from wheat, *ipt* from Agrobacterium, and *PLT5* from Arabidopsis^46–49^. For genome engineering, we provide sequences for eight different site-specific recombinases, each with a nuclear localization tag and codon-optimized for plant expression. Lastly, we provide domesticated elements for constructing geminiviral replicons based on Bean Yellow Dwarf Virus and Wheat Dwarf Virus. Modules are provided with the LIR-SIR-LIR (LSL) sequences for replicon initiation, with and without the replicon protein genes for construction of both *cis* and *trans* architectures for viral replication^50^. Each LSL module contains an Esp3I linker in the native position of the virion-sense gene products, compatible with the destination donor modules from the toolkit chassis to create L1 or L2 destination points. This method for replicon assembly is significantly more flexible than earlier methods, in which the T-DNA backbone could not be modified, or where replicon cargo was limited by modular capacity^26,31^. The complete Phytobrick library is enumerated in Supplementary Table 1; vector maps, curated with updated annotations, are found in Supplementary Fileset 1.

### Benchmarking *cis*-regulatory elements

#### Benchmarking dicot & monocot promoters

For *cis*-regulatory elements, we focused on cloning promoters and terminators from plant housekeeping genes with evidence in recent literature for effective transgene expression, especially for gene editing. Some have never been assayed for expression strength, or only by indirect readouts such as editing frequency, and few have been previously adapted to the Phytobrick standard. To quantitatively compare expression, we built constructs to test several panels of these elements with a firefly/Renilla dual-luciferase assay in infiltrated *N. benthamiana* leaves and *Setaria viridis* protoplasts. This high-throughput, ratiometric assay uses a fixed reporter to normalize for delivery between samples, providing a higher correlation with mRNA levels than single-reporter assays such as GFP intensity^11^. For the first panels, dicot promoters and terminators tested in leaf infiltrations, the construct backbone included Renilla driven by the *Nos* promoter and terminator; Ruby, to identify infiltration samples with good delivery; and a pair of enhancer blocking insulators, EXOB and TBS, included to minimize interference from other cassettes on the construct or any neighboring genomic elements (punches were collected at 5DAI, likely sampling expression from both transient and integrated T-DNAs). Between the insulators, we used an expansion module to place a Level 1 destination point for entry of a variable firefly cassette directly from Phytobrick parts (Figure 3a). All firefly cassettes for the dicot promoter panel used the *Nos* terminator, and those for the terminator panel used the *2x35S* promoter and *TMV*Ω 5’ leader sequence. The analogous construct backbone for the monocot promoter panel included additional marker genes intended to enable selection of stable transgenics, and substituted some regulatory elements targeted for monocot expression (Figure 3b).

**Figure 3:**
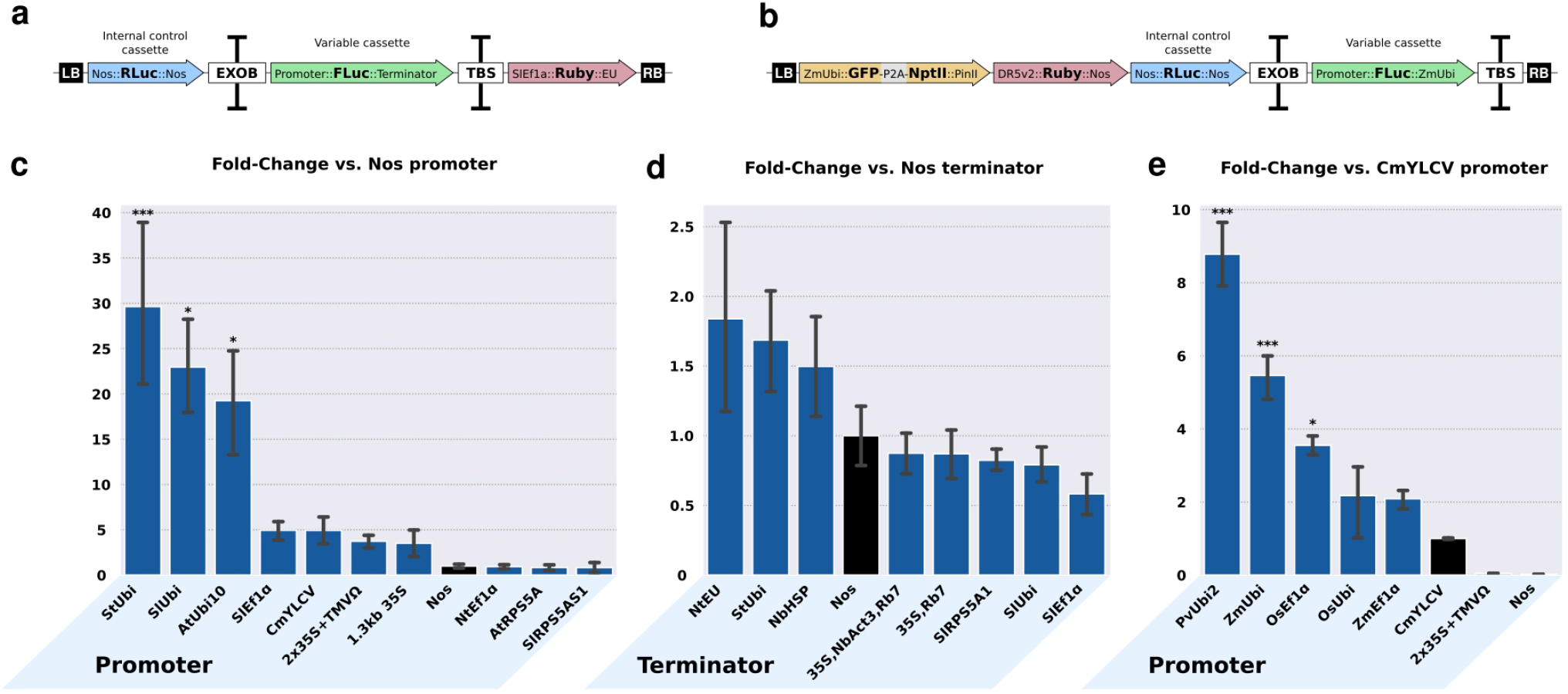
Benchmarking expression from promoter and terminator elements. **3a:** Vector archetype for dicot promoter and terminator test constructs. **3b:** Vector archetype for monocot promoter test constructs. **3c-e:** Results for promoter and terminator testing. Data are presented as the mean fold-change SE versus an internal control: *Nos∷Firefly∷Nos, 2x35S∷TMV*Ω*∷Firefly∷Nos*, and *CmYLCV∷Firefly∷ZmUbi* for the dicot promoter, dicot terminator, and monocot promoter panels respectively (control treatments highlighted in black). P-values between each construct and the control were calculated using pairwise *t*-tests and adjusted for multiple-testing: * indicates P *≤* 0.05, and *** indicates P *≤* 0.001.

The dicot promoter panel included orthologs of ubiquitin, Ef1*α*, and RPS5A from several species, the viral promoters *CmYLCV*, single and double *35S*, and a *Nos* control (Figure 3c). The Ef1*α* and RPS5A promoters were selected because certain orthologs have been shown to be highly effective drivers of CRISPR/Cas reagents for stable gene editing in Arabidopsis, Nicotiana, and tomato^8,51,52^. Those results were presumably due to strong expression of these promoters in meristematic or germline cells; nevertheless, it is useful to characterize their expression in infiltrated leaves, as different applications may benefit from both germline and somatic expression. In our assay, the ubiquitin promoters led to the highest expression, ~20-30 fold higher than *Nos*; the potato and tomato ubiquitin promoters were also significantly higher than *2x35S*. The *SlEf1α* promoter was moderately expressed, comparable to the viral promoters, while the *NtEf1α, SlRPS5A1* and *AtRPS5A* promoters exhibited the lowest expression, below the *Nos* control. Our results are generally consistent with previous reports comparing ubiquitin and viral promoters in dicots^5,11^, and with available RNA-seq data for the corresponding loci in tomato and Arabidopsis. However, they also underscore the importance of experimentally validating the performance of regulatory elements, especially across species. For example, the *AtUbi10* promoter performed well in our *N. benthamiana* system, while the more closely related *NtEf1α* promoter from tobacco was relatively poor, despite moderate to high predicted expression of the orthologous *SlEf1α* promoter in tomato. Differences could be due to evolutionary distance, but also the particular sequences cloned; previous study indicates promoter sequence length matters, presumably due to variable inclusion of upstream regulatory sequences^5^. All of our newly cloned promoters, inclusive of large 5’ introns, were kept ≤ 1.7kb for ease of cloning, and with the exception of *NtEf1α*, these retained predicted expression in leaf tissue. Between our research and others, the *SlEf1α* promoter has now shown evidence of strong expression in both somatic and germline tissues, and may represent a good default choice for gene editing experiments in Solanaceous species.

The monocot promoter panel compared the same classic promoters to cereal orthologs of ubiquitin and Ef1*α* (Figure 3e). Similar to the dicot panel, we saw strongest expression from ubiquitin promoters, *PvUbi2* and *ZmUbi*, followed by the *OsEf1α* promoter. All three were significantly higher than the *CmYLCV* control. The *OsUbi* and *ZmEf1α* promoters were also higher than *CmYLCV*, though these differences were not significant. Both the *2x35S* and *Nos* promoters performed poorly. Weak expression from *Nos* was particularly notable: we observed low Renilla expression across all samples, independent of the firefly measurement. Indeed, we originally attempted to measure luciferase activity in transgenic rice callus lines; however, we found that with slightly higher levels of background signal in wild-type callus controls, Renilla expression in many events was too low to clearly discern a ratiometric readout, leading us to test the same constructs instead with an efficient setaria protoplast system^53^. To the best of our knowledge, no previous promoter comparison in monocots has included *Nos, CmYLCV, OsUbi, OsEf1α* or *ZmEf1α*. The performance of *Nos* may be explained by evolutionary origin, as monocots are not among the natural host range of Agrobacterium. Ours is also not the first study to find *ZmUbi* outperforms *35S* in cereals; previous studies found fold-change differences varying from less than 10 to close to 100^54–58^, and our data are consistent with the higher end of this range. For the newly tested housekeeping gene promoters, performance was mixed with respect to predictions from RNA seq data in maize and rice. In both species, ubiquitin is more highly expressed than Ef1*α* in leaf tissue, while *ZmEf1α* is more highly expressed than *ZmUbi* in maize embryos, and *OsEf1α* than *OsUbi* in rice callus. The maize promoters were consistent with this expectation in mesophyll-derived setaria protoplasts, but the inverse was true for the rice promoters. As with the dicot promoters, these results could be attributable to evolutionary distance (setaria is more closely related to maize than rice) or to particulars of the cloned sequences.

#### Benchmarking dicot terminators

Many previous studies have indicated an important role for 3’ regulatory regions in transgene expression, though only a few have systematically compared large panels of 3’ UTR/terminator regions^10,11^. Mechanistically, proper transcriptional termination is understood as essential to minimizing post-transcriptional gene silencing (PTGS), effected by RDR6-mediated targeting of unpolyadenylated transcripts that become enriched during transcriptional read-through^59,60^. Compound terminators, which stack two or more single terminators together, have been found to increase transgene expression in both transient and stable expression systems, though to widely varying degrees^10,59,61^. We assembled a terminator panel consisting of new 3’ regions corresponding to several previously tested promoters (*StUbi, SlUbi, SlEf1α*, and *SlRPS5A1*), and a subset of terminators previously tested by Diamos & Mason (2018) spanning their observed range of performance (Figure 3d). These previously tested terminators include *Nos*, among the weakest, and both medium- and high-performing single and compound terminators *NtEU, NbHSP, 35S+Rb7*, and *35S+NbAct3+Rb7*. The latter two feature a Rb7 matrix attachment region (MAR), found to increase gene expression when placed 3’ to multiple terminators^10^. Surprisingly, we observed only modest differences in expression, none of which were statistically significant. The difference between *Nos* and the highest-expressing terminator, *NtEU*, was less than 2-fold; by contrast, the previous study found greater than 10-fold difference, using the same promoter, leaf infiltration system, and time to sample collection. One possible explanation is use of different reporters, as they quantified intensity of a single GFP marker. Besides normalizing for variable delivery, our dual-luciferase system may also produce different results as a function of protein half-lives, which are far shorter for both firefly and Renilla than fluorescent proteins such as GFP^62^. By way of comparison, the group-wide fold-change differences we observed were more in line with those observed by Tian *et al.* (2022)^11^, who also used a dual-luciferase system. Unfortunately, the dearth of overlapping elements among our respective panels makes it difficult to conclude if these results are intrinsic reflections of the terminators themselves or a true technical effect of the assay. The majority of previous studies of terminator function in both transient and stable transgenic systems, comparing different single or double terminators, have found ~2-5 fold differences between treatments, suggesting a real effect on expression which is nevertheless modest in comparison to promoter selection^61,63,64^. Given these divergent results, further research seems warranted to better elucidate the role of terminators on expression.

#### Screening promoters for enhancer activity

Lastly, we undertook a study of enhancer activity among the highly-expressed promoters from our dicot panel. Enhancers are *cis*-regulatory elements that increase gene expression by one or more mechanisms, generally involving transcription factor recruitment or chromatin interactions^65^. As with expression from defined promoter sequences, enhancer function may be constitutive or tissue-specific, but unlike proximal promoter elements, they may be located multiple kilobases away from coding sequences, and act in orientation-independent fashion^12,66^. More than a dozen enhancer sequences have been identified in plants, but few have been interrogated in a biotechnology context^67^. The sole well-studied example is the near-upstream enhancer of the *35S* promoter, with greatest activity in the *2x35S* variant. Nearly all generalized features of plant enhancers are based on studies using the *35* enhancer to activate various tissue-specific promoters: these characteristics include bidirectional function, and degree dependence on both physical distance and the specific secondary promoter subjects^42,68,69^. Trans-activation of nearby promoters is a particular concern for coordinated transgene expression, as enhancer activity can abrogate the specificity of inducible or tissue specific promoters. One solution is to place an insulator sequence between the enhancer and subject promoter, which can fully or partially rescue specific expression^13^. However, this requires an understanding of those promoters containing trans-acting enhancers, and the few reports investigating off-target activity of promoters besides *35S* are inconclusive as to whether this is a rare or general feature of strong, constitutive promoters. One found an effect with *35S* but not with *Nos*, and another found comparable effects from *35S* and a *Nos/Mas* chimeric super-promoter, diminished effects from a rubisco promoter or *Nos* alone, and no effect from a tobacco cryptic promoter^70,71^. Lack of study of promoter enhancer activity therefore represents a serious knowledge gap.

To assess potential enhancer activity, we used a modified dual-luciferase assay with each of two weak promoters: *GmHSP*, a heat-shock inducible promoter, and *GmGy1*, a seed-specific promoter, neither of which exhibit appreciable expression in our leaf infiltration testbed. Each construct contained fixed Ruby and Renilla markers with an intervening insulator sequence, as in earlier experiments, while the variable promoters with potential enhancer activity drove an unused selectable marker placed head-to-head with a firefly cassette driven by either weak promoter (Figure 4a). We tested each of the promoters from the dicot panel, excluding the lowly-expressed RPS5A orthologs, against both weak promoters, and looked for increased firefly expression relative to Renilla (Figure 4b). The negative control was a promoterless marker cassette. For each of these samples, firefly signal was below the limit of detection (determined by background signal in WT samples), while Renilla signal was always present; this is consistent with expectation that, in the absence of enhancer activity, the weak promoters are not expressed in leaf tissue. Interestingly, the only treatments exhibiting an observable enhancer effect were the viral promoters: both *2x35S* and *CmYLCV* led to significant activation of *GmGy1*. In the corresponding *GmHSP* treatments, some samples exhibited detectable firefly expression, but none showed relative expression significantly higher than the negative control. None of the housekeeping gene promoters exhibited a significant activation effect on either weak promoter, and most samples were below the limit of detection.

**Figure 4:**
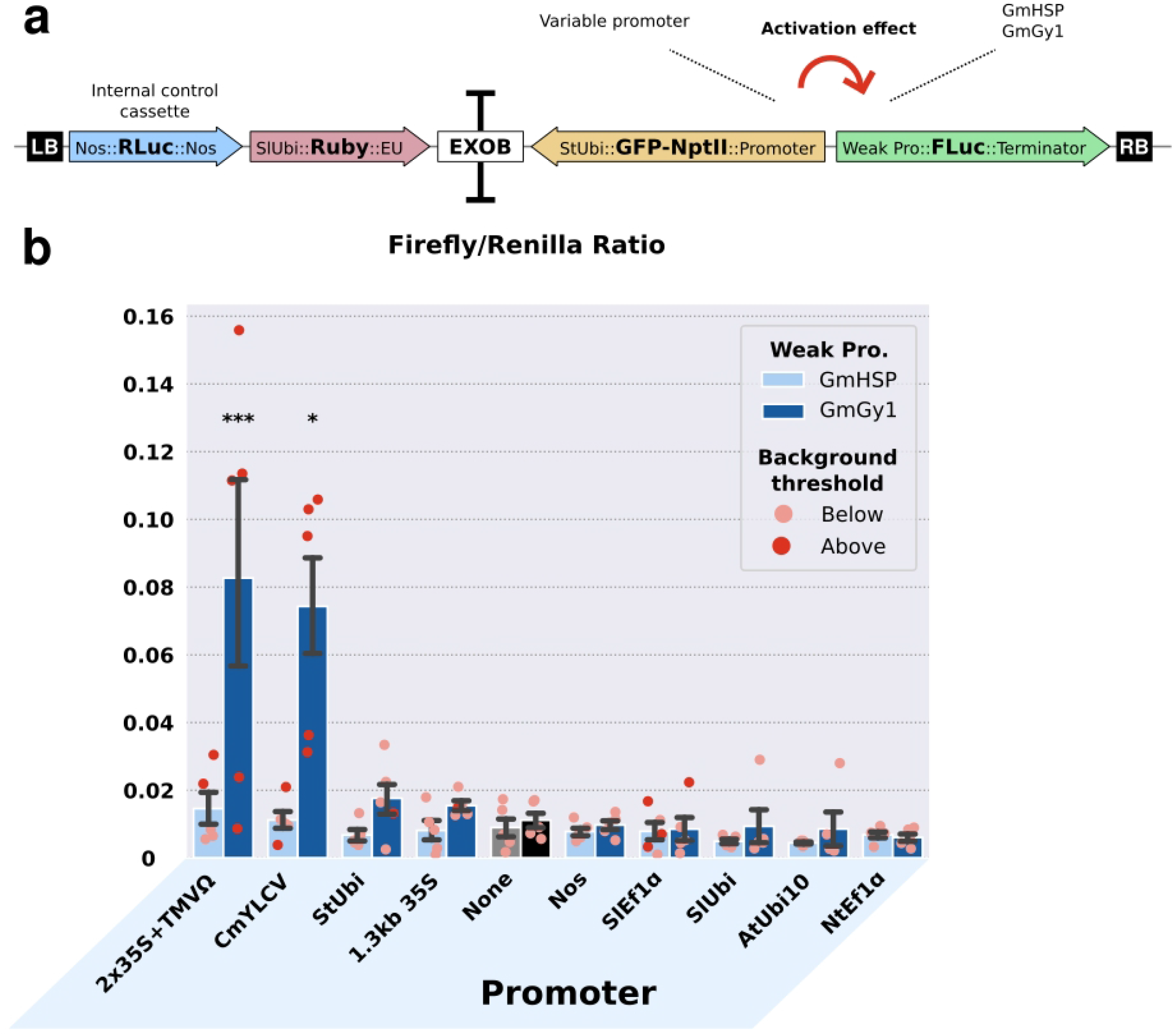
Screening promoters for enhancer activity. **4a.** Vector archetype for promoter trans-activation test constructs. **4b.** Results for promoter trans-activation testing. Data are presented as the mean firefly/Renilla ratio ± SE. Constructs with the GmHSP promoter are shown in light blue, and those with the GmGy1 promoter are shown in dark blue. Individual sample values are superimposed on bar charts. Dark red points indicate samples with firefly values above the experimentally determined background threshold of 150 RLUs; light red points indicate samples with firefly values below this threshold, which are effectively indistinguishable from zero. MP-values between each construct and the control treatments (with no promoter, highlighted in grey/black) were calculated using pairwise *t*-tests and adjusted for multiple testing: * indicates P *≤* 0.05, and *** indicates P *≤* 0.001.

Several points are notable from our results. First, consistent with earlier literature, we found that activation depends on the subject promoter, as we observed activation of *GmGy1* but not *GmHSP* from the same enhancers. Second, it appears there is no correlation between on-target and off-target expression, as the ubiquitin promoters drove 4-6 fold greater expression than the viral promoters in the on-target promoter assay, yet led to no detectable trans-activation in this assay. Lastly, while we certainly cannot rule out all potential for enhancer activity, our results suggest this may not a general feature of housekeeping gene promoters; conversely, viral promoters may be more likely to carry constitutive enhancers. In any case, housekeeping gene promoters seem a safer choice for use in constructs where specific transgene expression is required.

## Conclusion

Our overarching goal in this paper was to increase the scope of community access to genetic resources for plant engineering. We accomplished this through cloning designs that 1) emphasize compatibility with previous reagents, such as the Phytobrick library and binary vectors, and 2) streamline the assembly steps required to build complex constructs. This allows researchers to focus on construct design over implementation, especially beneficial for labs without extensive experience in molecular cloning. Construct design is further guided by the results of our *cis*-regulatory element experiments, revealing patterns of transgene expression in multiple systems. The vectors are available for research use through two separate collections on Addgene: the Phytobrick library (plate ID *XXXXXXXX*), and the JMC toolkit (plate ID *YYYYYYYY*). The supplement includes step-by-step protocols for each type of Golden Gate assembly, and troubleshooting tips for complex assemblies. In summary, our work will empower researchers to better leverage both current and emerging tools for diverse applications in plant research.

## Supporting information

Supplementary Fileset 1

Supplementary Fileset 2

Supplementary Protocols

## Acknowledgements

This work was supported by the funding from the National Science Foundation (NSF-IOS-20402180) and the U.S. Department of Energy (DE-SC0018277) to D.F.V. J.K. and F.Z. were partially supported by USDA-NIFA (2021-67013-34565) and NSF PGRP (IOS-2040218) awards.

## Supplementary Materials

Supplementary Figures: Toolkit illustrations

Supplementary Table 1: Toolkit vectors database spreadsheet

Supplementary Fileset 1: Toolkit vector maps

Supplementary Fileset 2: Experiment construct vector maps

Supplementary Protocols: Cloning protocols

