## Supplementary Figures for "An Extensible Vector Toolkit and Parts Library for Advanced Engineering of Plant Genomes"

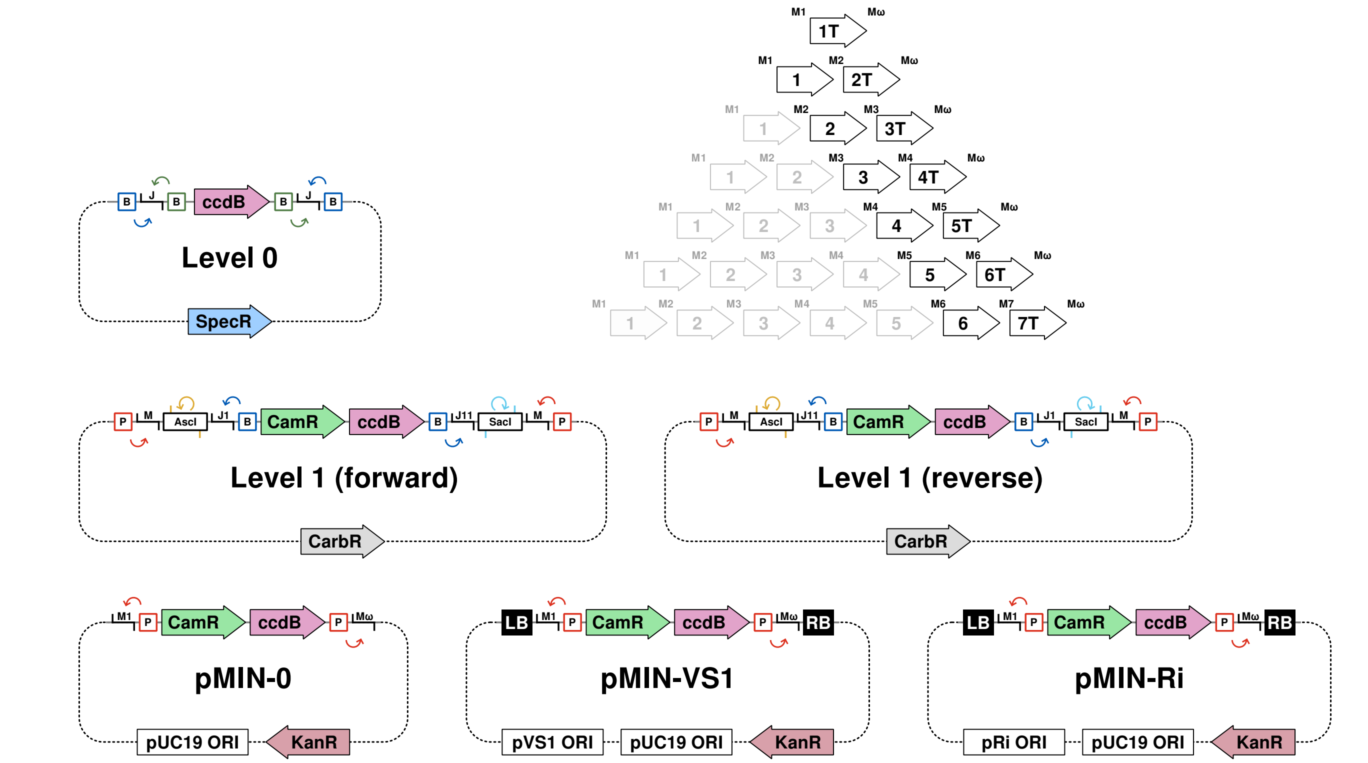


**Figure S1: Destination vectors for hierarchical assembly.** Destination vectors, from Level 0 through Level 2, use *ccdB* counterselection. Level 1 destination vectors are available for each position in the 7-module circuit with both forward and reverse orientation Phytobrick entry, and feature AscI and SacI sites for traditional cloning in addition to BsaI Golden Gate entry.

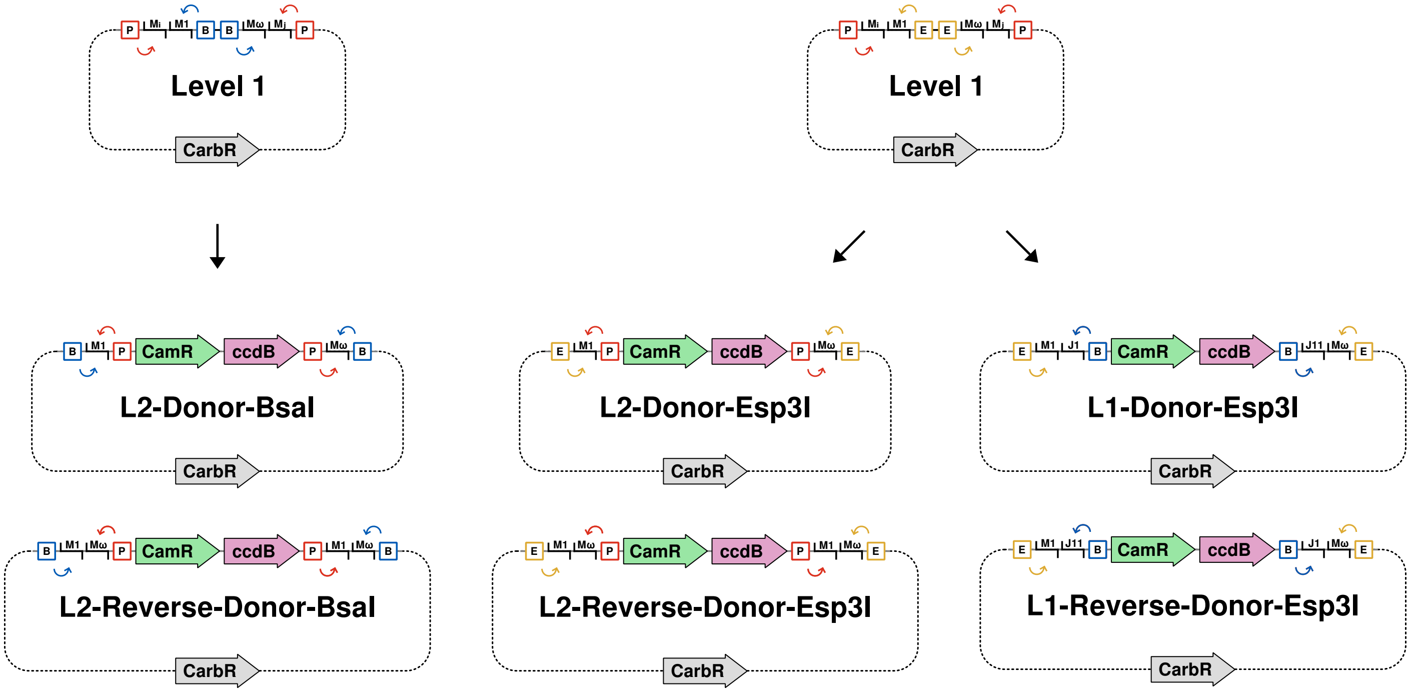


**Figure S2: Vector sets for modular expansion system.** Level 1 expansion modules contain internal Esp3I or BsaI linkers. BsaI linkers can be used to introduce L2 donors, and Esp3I linkers can be used to introduce L1 or L2 donors.
