## Supplementary Protocols for "An Extensible Vector Toolkit and Parts Library for Advanced Engineering of Plant Genomes"

### Supplementary Protocol 1: Level 0 (Phytobrick) Assembly

1. Design a PCR product or synthetic fragment with BbsI sites creating Phytobrick overhangs, flanking the desired functional unit and matching one of the compatible destination vectors: **pMC-1-ccdB-6** (promoter), **pMC-1-ccdB-4** (promoter with room for inclusion of additional 5’ UTR or recombinase recognition sequence), **pMC-6-ccdB-9** (coding sequence), or **pMC-9-ccdB-11** (terminator), **pMC-1-ccdB-11** (full-length unit, e.g. for non-coding components such as insulators). For detailed instructions on Phytobrick design and domestication strategies, see Cai, Lopez & Patron (2020)(Cai et al., 2020).
2. Set up a Golden Gate reaction and run with the following thermocycler program: 5x (37${}^{\circ}$C/5min + 16${}^{\circ}$C/10min) + 37${}^{\circ}$C/15min + 80${}^{\circ}$C/5min

- 2$\mu$L T4 DNA Ligase Buffer
- 1$\mu$L T4 DNA Ligase
- 0.5$\mu$L BbsI-HF (NEB R3539)
- 40ng Level 0 Destination Vector
- 20-40ng Insert (amplicon or fragment)
- $H_{2}0$ to 20$\mu$L

3. Transform 5$\mu$L of the assembly mix into NEB10$\beta$. Plate on LB with 50mg/L spectinomycin and grow overnight at 37${}^{\circ}$C.
4. Pick 2-3 colonies into 5mL liquid cultures, grow for 16-18h overnight at 37${}^{\circ}$C, then miniprep.
5. Confirmation: typically, >95% of clones will feature the correct insert. However, as both PCR products and synthetic fragments are subject to occasional SNPs, we recommend sequencing the entire insert to ensure a perfect clone. The following primers anneal to all Level 0 backbones, and typically provide complete coverage in Sanger sequencing for inserts $\leq$ 1.3kb:

- Forward: 5’ GCCTTTGAGTGAGCTGATACC 3’
- Reverse: 5’ GTCATGATAATAATGGTTTCTTAGACGTCA 3’

##### Notes

- The *ccdB* destination vectors provided with our toolkit are intended for simplified Phytobrick classes. For assembly of the more atomic Phytobrick classes, *lacZ*$\alpha$ destination vectors are available from the MoClo Plant Parts Kit, Addgene Kit #1000000047(Engler et al., 2014).
- Our lab uses BbsI-HF. However, the vectors and reaction are also compatible with the isoschizomer BpiI (Thermo Scientific ER1011).

### Supplementary Protocol 2A: Level 1 Assembly from Phytobricks

1. Select a set of Level 0 parts forming a contiguous Phytobrick unit, e.g. a single promoter (J1-J6), coding sequence (J6-J9), and terminator (J9-J11), or a single full-length part (J1-J11). Select any Level 1 destination vector: **pJMC-1T** through **pJMC-7T** for forward-orientation cassettes, or **pJMC-1T-Reverse** through **pJMC-7T-Reverse** for reverse-orientation cassettes.
2. Set up a Golden Gate reaction and run with the following thermocycler program: 10x (37${}^{\circ}$C/5min + 16${}^{\circ}$C/10min) + 37${}^{\circ}$C/15min + 80${}^{\circ}$C/5min

- 2$\mu$L T4 DNA Ligase Buffer
- 1$\mu$L T4 DNA Ligase
- 0.5$\mu$L BsaI-HFv2 (NEB R3733)
- 50ng Level 1 Destination Vector
- 40ng Each Insert
- $H_{2}0$ to 20$\mu$L total

3. Transform 5$\mu$L of the assembly mix into NEB10$\beta$. Plate on LB with 50mg/L carbenicillin and grow overnight at 37${}^{\circ}$C.
4. Pick 2 colonies into 5mL liquid cultures, grow for 16-18h overnight at 37${}^{\circ}$C, then miniprep.
5. Confirmation: typically, >95% of clones will feature the correct assembly. Since reaction components are (presumably) sequenced plasmids, sequencing of the assembly junctions is sufficient to ensure clone integrity. The following primers anneal to all Level 1 backbones, and provide coverage in Sanger sequencing for the terminal junctions J1 and J11:

- Forward: 5’ GCCTTTGAGTGAGCTGATACC 3’ (same primer as Level 0 backbone)
- Reverse: 5’ GTATCCAACATTTCCGTGTCGC 3’

##### Notes

- NEB has discontinued earlier versions of BsaI. We specifically recommend use of the current BsaI-HFv2, which is optimized for Golden Gate assembly and activity in T4 ligase buffer.

### Supplementary Protocol 2B: Level 1 Assembly with U6 sgRNA Cassette

Overview: this protocol offers a simple way to assemble single guide RNA (sgRNA) cassettes for SpCas9 with either AtU6 or OsU6 promoters, using annealed oligos to introduce the spacer. The protocol uses BsaI and terminal overhangs J1 and J11, for assembly into any Level 1 module. However, unlike a standard Phytobrick assembly, it uses custom internal overhangs to permit assembly of the variable spacer sequence. Reagents for Phytobrick-compatible assembly of more complex CRISPR targeting reagents, such as tRNA arrays, are available from the MoClo CRISPR/Cas Toolkit for Plants, Addgene Kit #1000000159(Hahn et al., 2020). The advantage of this protocol is that Level 1 assembly is accomplished in a single cloning step (rather than first subcloning the spacer into a Level 0 destination vector).

1. Select any Level 1 destination vector: **pJMC-1T** through **pJMC-7T** for forward-orientation cassettes, or **pJMC-1T-Reverse** through **pJMC-7T-Reverse** for reverse-orientation cassettes. For the inserts, select either **pMC-1-AtU6-Y** or **pMC-1-OsU6-X** as the promoter; and use **pMC-Z-SpCas9-gRNA-repeat-polyT-11**.
2. Synthesize a pair of DNA oligos to introduce the 20 spacer sequence, following the design pattern below for each promoter. The 5’ end of the forward oligo provides a 4bp overhang (5’ $\to$ 3’ GATT or TTGT) specific to the selected promoter, while the 5’ end of the reverse oligo provides a 4bp overhang matching the scaffold (5’ $\to$ 3’ AAAC). Replace the N bases with the desired 20bp spacer and its reverse complement, respectively. Note that the first nucleotide for U6 transcription must be a G; if the spacer also starts with a G, the length of variable N sequence may be reduced to 19.

```
AtU6: GATTGNNNNNNNNNNNNNNNNNNNN
          |||||||||||||||||||||

          CNNNNNNNNNNNNNNNNNNNNCAAA
OsU6: TTGTGNNNNNNNNNNNNNNNNNNNN
          |||||||||||||||||||||

          CNNNNNNNNNNNNNNNNNNNNCAAA
```

3. Phosphorylate and anneal the oligo pair. Set up a reaction and run with the following thermocycler program: 37${}^{\circ}$C/30min + 98${}^{\circ}$C/1min, followed by a ramp down from 98${}^{\circ}$C to 25${}^{\circ}$C at 1${}^{\circ}$C/second. Once done, prepare a 25x dilution of the reaction.

- 3$\mu$L T4 DNA Ligase Buffer
- 2$\mu$L T4 Polynucleotide Kinase (NEB M0201)
- 3$\mu$L Forward Oligo @ 100$\mu$M
- 3$\mu$L Reverse Oligo @ 100$\mu$M
- $H_{2}0$ to 30$\mu$L total

4. Set up a Golden Gate reaction and run with the following thermocycler program: 10x (37${}^{\circ}$C/5min + 16${}^{\circ}$C/10min) + 37${}^{\circ}$C/15min + 80${}^{\circ}$C/5min

- 2$\mu$L T4 DNA Ligase Buffer
- 1$\mu$L T4 DNA Ligase
- 0.5$\mu$L BsaI-HFv2 (NEB R3733)
- 50ng Level 1 Destination Vector
- 40ng **pMC-1-AtU6-Y** OR **pMC-1-OsU6-X**
- 40ng **pMC-Z-SpCas9-gRNA-repeat-polyT-11**
- 1$\mu$L 25x Diluted Oligo Linker
- $H_{2}0$ to 20$\mu$L total

5. Transform 5$\mu$L of the assembly mix into NEB10$\beta$. Plate on LB with 50mg/L carbenicillin and grow overnight at 37${}^{\circ}$C.
6. Pick 2 colonies into 5mL liquid cultures, grow for 16-18h overnight at 37${}^{\circ}$C, then miniprep.
7. Confirmation: typically, >95% of clones will feature the correct assembly. Given the short length of a U6 cassette, sequencing of the entire insert can be achieved with the same primers as a standard Phytobrick Level 1 assembly (Protocol 2A).

### Supplementary Protocol 3: Level 2 Assembly

1. Select a set of Level 1 modules forming a complete chain (see Figure S1). Select a Level 2 destination vector: **pMIN-0**, **pMIN-VS1**, **pMIN-Ri**, or any Level 2 destination vector previously assembled through the vector extension toolkit.
2. Set up a Golden Gate reaction and run with the following thermocycler program: 10x (37${}^{\circ}$C/5min + 16${}^{\circ}$C/10min) + 37${}^{\circ}$C/15min + 80${}^{\circ}$C/5min

- 2$\mu$L T4 DNA Ligase Buffer
- 1$\mu$L T4 DNA Ligase
- 0.5$\mu$L PaqCI (NEB R0745)
- 0.4$\mu$L PaqCI Activator (included with NEB R0745)
- 60-80ng Level 2 Destination vector
- 50-80ng Each Level 1 Module
- $H_{2}0$ to 20$\mu$L total

3. Transform 5$\mu$L of the assembly mix into NEB10$\beta$. Recover in 100-200$\mu$L SOC for 1 hour at 37${}^{\circ}$C, then plate on LB with 50mg/L kanamycin and grow overnight at 37${}^{\circ}$C.
4. Pick 2-6 colonies into 5mL liquid cultures, grow for 16-18h overnight at 37${}^{\circ}$C, then miniprep.
5. Confirmation: typically, a high fraction of clones are correct. For simpler assemblies, with 3-4 parts, the fraction is >95%. For more complex assemblies, with up to 7 inserts, larger or more variable sized modules, or very large backbones, the fraction may decrease to ~60-80%. For this reason, we recommend screening a small number of clones by digest prior to sequencing. AscI and SacI are often good diagnostic enzymes, as they appear once in each Level 1 module insert. Final confirmation may be achieved by Sanger sequencing of assembly junctions. For larger assemblies, whole-plasmid sequencing is an economical alternative; we regularly confirm constructs using the Oxford Nanopore service from Plasmidsaurus (Eugene, OR, USA)

##### Notes

- Our lab uses PaqCI. However, the vectors and reaction are also compatible with the isoschizomer AarI (Thermo Scientific ER1581).
- For complex assemblies, efficiency can be greatly increased by substitution of high-concentration T4 DNA ligase (NEB M0202T), and by increasing the number of Golden Gate cycles.

### Supplementary Protocol 4: Extension Assembly via BsaI or Esp3I

1. Assemble a Level 2 construct including one of the expansion destination modules: **pJMC-1-BsaI** through **pJMC-7T-BsaI**, or **pJMC-1-Esp3I** through **pJMC-7T-Esp3I**. Select a destination donor module: for BsaI expansion, compatible modules are **pJMC-L2-Donor-BsaI** or **pJMC-L2-Reverse-Donor-BsaI** to create a new Level 2 destination point; for Esp3I expansion, compatible modules are **pJMC-L2-Donor-Esp3I** or **pJMC-L2-Reverse-Donor-Esp3I** to create a new Level 2 destination point, or **pJMC-L1-Donor-Esp3I** or **pJMC-L1-Reverse-Donor-Esp3I** to create a Level 1 destination point.
2. Set up a Golden Gate reaction and run with the following thermocycler program: 5x (37${}^{\circ}$C/5min + 16${}^{\circ}$C/10min) + 37${}^{\circ}$C/15min + 80${}^{\circ}$C/5min

- 2$\mu$L T4 DNA Ligase Buffer
- 1$\mu$L T4 DNA Ligase
- 0.5$\mu$L BsaI-HFv2 (NEB R3733) OR Esp3I (NEB R0734)
- 60-80ng Level 2 construct
- 60ng Destination donor module
- $H_{2}0$ to 20$\mu$L total

3. Transform 5$\mu$L of the assembly mix into DB3.1. Recover in 100-200$\mu$L SOC for 1 hour at 37${}^{\circ}$C, then plate on LB with 50mg/L kanamycin and 25mg/L chloramphenicol and grow overnight at 37${}^{\circ}$C.
4. Pick 1 colony into a 5mL liquid culture, grow for 16-18h overnight at 37${}^{\circ}$C, then miniprep.
5. Confirmation: with double antibiotic selection, clones are nearly universally correct (>99%). Sequencing confirmation of the assembly junctions can be achieved using the following primers, which anneal to all the destination donor insert modules:

- Forward: 5’ GTGATATTATTGACACGCCC 3’
- Reverse: 5’ GTACCTATAACCAGACCGTTCA 3’

##### Notes

- Our lab uses Esp3I. However, the vectors and reaction are also compatible with the isoschizomer BsmBI-v2 (NEB R0739). Both enzymes function in T4 ligase buffer, but BsmBI-v2 is most active at 42${}^{\circ}$C and may benefit from a modified thermocycling program. See NEB E1601 for details.

### Tips & Troubleshooting Steps

- The reactions, particularly simpler assemblies such as Level 0, Level 1, and extension reactions, tolerate a range of input DNA masses. Our mass recommendations are based on a target of 20 fmol for each DNA component, following the most recent protocol from the Marillonet group(Marillonnet & Grützner, 2020). For complex assemblies, calculating input mass on a part-by-part basis may increase efficiency. The input volume in $\mu$L for a component is given by the formula: [20 fmol * length (bp)] / [1520 * concentration (ng/$\mu$L) ].
- Reaction efficiency varies with the quality of the input DNA. We have found DNA dilutions from midipreps produce higher reaction efficiency than minipreps, and a failing assembly may often be rescued by substituting a fresh preparation or dilution, particularly for the destination vector.
- The number of recommended digestion/ligation cycles for each assembly is based on our experience with typical reaction efficiencies, in terms of the number of clones recovered following transformation into *E. coli*. For low-efficiency reactions, the number of clones can be greatly increased by increasing the number of cycles to 20 or even 50 cycles (run as an overnight program). This is typically only required for very complex assemblies.
- Cloning efficiency is also greatly affected by the selection and preparation of competent cells for transformation. We typically use chemically competent NEB10$\beta$ cells, but have also had success using NEB Stable cells, DH5$\alpha$, and similar *ccdB*-sensitive strains. New competent cell preparations should be screened for efficiency using a confirmed plasmid, and screened for background contamination on the relevant antibiotics. This is particularly important for bacteriostatic antibiotics such as carbenicillin which may permit formation of satellite colonies.
